# Thrombopoietin from hepatocytes promotes hematopoietic stem cell regeneration after myeloablation

**DOI:** 10.1101/2021.05.13.443980

**Authors:** Longfei Gao, Matthew Decker, Haidee Chen, Lei Ding

**Author notes:** These authors contribute equally (Longfei Gao, Matthew Decker). Author for correspondence: Columbia Stem Cell Initiative, Department of Rehabilitation and Regenerative Medicine, Department of Microbiology and Immunology, Columbia University Irving Medical Center, 650 West 168^th^ ST, BB 1103C, New York, NY, 10032.

## Abstract

The bone marrow niche plays a critical role in hematopoietic recovery and hematopoietic stem cell (HSC) regeneration after myeloablation. However, it is not clear whether systemic factors beyond the local niche are required for these essential processes *in vivo*. Thrombopoietin (TPO) is a critical cytokine promoting hematopoietic rebound after myeloablation and its transcripts are expressed by multiple cellular sources. The upregulation of bone marrow-derived TPO has been proposed to be crucial for hematopoietic recovery and HSC regeneration after stress. Nonetheless, the cellular source of TPO in stress has never been investigated genetically. We assessed the functional sources of TPO following two common myeloablative perturbations: 5-fluorouracil (5-FU) administration and irradiation. Using a *Tpo* translational reporter, we found that the liver but not the bone marrow is the major source of TPO protein after myeloablation. Mice with conditional *Tpo* deletion from osteoblasts or bone marrow stromal cells showed normal recovery of HSCs and hematopoiesis after myeloablation. In contrast, mice with conditional *Tpo* deletion from hepatocytes showed significant defects in HSC regeneration and hematopoietic rebound after myeloablation. Thus, systemic TPO from the liver is necessary for HSC regeneration and hematopoietic recovery in myeloablative stress conditions.

## Introduction

Hematopoietic stem cells (HSCs) generate all blood and immune cells, and play critical roles in the regeneration of the blood system after stress. They reside in the bone marrow niche where Leptin Receptor^+^ (LepR^+^) perivascular stromal cells and endothelial cells are the major sources of factors that promote their maintenance ^1–3^. Following myeloablative stress, HSCs regenerate in the bone marrow niche, which is essential for the recovery of hematopoiesis ^4–7^. Several signaling pathways, including Notch, EGF/EGFR, pleiotrophin, Dkk1, Angiogenin, and SCF/c-KIT, have been shown to be critical for this process ^8–13^. Many of these cytokines arise from the local bone marrow niche for HSC regeneration and hematopoietic recovery in a paracrine fashion ^12,14-16^. However, it is not clear whether systemic signals that originate outside the bone marrow also regulate HSC regeneration and hematopoietic recovery after stress.

Thrombopoietin (TPO) is critical for bone marrow HSC maintenance and hematopoietic recovery ^17–20^. TPO was first identified as the ligand to the MPL receptor and a driver of megakaryopoiesis and platelet production ^21–23^. It was later shown that the TPO/MPL signaling was required for the maintenance of HSCs ^24^. The TPO/MPL signaling also plays a crucial role in hematopoietic stress response, particularly after myeloablation. The characteristic hematopoietic progenitor rebound following administration of the antimetabolite drug 5-fluorouracil (5-FU) is dependent on MPL ^25^. After irradiation, the TPO/MPL signaling is similarly essential for hematopoietic recovery and survival ^20,26-28^. Indeed, TPO mimetic drugs, such as romiplostim and eltrombopag, have been shown to improve recovery after ablative challenge, and have also been used clinically to support hematopoiesis in diseases such as immune thrombocytopenic purpura and aplastic anemia ^29–32^.

The regulation of TPO production has been extensively investigated, but the *in vivo* source of TPO for HSC regeneration and hematopoietic recovery after myeloablation is not clear.

Previous studies have found that bone marrow populations such as stromal cells and osteoblasts may upregulate TPO in hematopoietic stress conditions, while the liver produces *Tpo* transcripts at a constant level ^19,33^. However, other investigators have found no significant changes in bone marrow *Tpo* transcript levels after 5-FU-mediated myeloablative treatment ^25^. Because *Tpo* expression is under heavy translational control ^34^, it is not clear what cells produce TPO protein for HSC regeneration and hematopoietic recovery after myeloablation. Furthermore, although upregulation of TPO may be a key mechanism of the bone marrow response to hematopoietic stress, the role of local TPO from bone marrow niche and systemic TPO from the liver for HSC and hematopoietic recovery has not been functionally investigated *in vivo*. Nonetheless, most studies proposed that local TPO derived from the bone marrow niche is critical for HSC and hematopoietic recovery after myeloablation ^19,35^. Here we genetically dissected the *in vivo* source of TPO for HSC and hematopoietic recovery following myeloablative stress.

## Results

### Myeloablation induced by 5-FU drives TPO-dependent hematopoietic recovery and HSC expansion

5-FU is a commonly used chemotherapy agent that leads to myeloablation. To test whether TPO is required for hematopoietic recovery after 5-FU treatment, we administrated 5-FU to *Tpo* knockout (*Tpo^gfp/gfp^*) ^17^ and wild-type control mice at 150mg/kg *via* a single intraperitoneal injection. Because *Tpo^gfp/gfp^* mice may have phenotypes compared with wild-type controls without any treatment ^17^, we also normalized hematopoietic parameters with baseline mice of the same genotype without any treatment. Ten days after the 5-FU administration, treated wildtype mice showed normal leukocyte and neutrophil counts, normal reticulocyte frequency, but significantly increased platelet counts, while treated *Tpo^gfp/gfp^* mice had no platelet expansion (Fig. S1 A-D). When normalized to baseline levels, *Tpo^gfp/gfp^* mice had a significant reduction of leukocyte, neutrophil, and platelet counts as well as reticulocyte frequency compared with wildtype controls (Fig. S1 A-D), suggesting that *Tpo* is required for hematopoietic recovery. Normalized post-treatment bone marrow cellularity did not differ significantly while spleen cellularity was significantly reduced in *Tpo^gfp/gfp^* mice compared with wild-type controls (Fig. S1 E and F). The frequencies of bone marrow and spleen HSCs (Lineage^-^ Sca-1^+^ c-kit^+^ CD150^+^CD48^-^, LSKCD150^+^CD48^-^) were more than 10 fold greater in 5-FU treated wild-type mice than baseline controls, while 5-FU treated *Tpo^gfp/gfp^* mice showed no change from baseline (Fig. S1 G and H). Overall, 5-FU stimulated a 3.7-fold increase of total HSC number in wild-type mice, but no significant effects were observed in *Tpo^gfp/gfp^* mice (Fig. S1 I). Compared with wild-type controls, the HSC increase responsiveness to 5-FU in *Tpo^gfp/gfp^* mice was 8 fold less (Fig. S1 I). The effects of 5-FU on bone marrow and spleen hematopoietic progenitor cells (LSK) were also blunted in *Tpo^gfp/gfp^* mice (Fig. S1 J and K). Thus, hematopoietic recovery, expansion of HSCs and hematopoietic progenitors following chemoablation mediated by 5-FU is TPO-dependent.

### Hepatic but not bone marrow TPO is required for HSC and hematopoietic recovery after 5-FU-mediated myeloablation

To identify the source of TPO, we investigated *Tpo* expression following 5-FU treatment. It has been reported that the bone marrow upregulates *Tpo* mRNA in stress conditions ^33^. Although bone marrow stromal cells (CD45^-^ Ter119^-^) express *Tpo* transcripts, we did not observe a significant upregulation of *Tpo* transcripts 14 days after 5-FU injection compared with baseline levels (Fig. S2 A). TPO protein production is under strong translational controls ^34^. To assess translation and generation of TPO protein following 5-FU treatment, *Tpo^creER^; ZsGreen^LSL^* translational reporter mice ^17^ were injected with 5-FU and immediately started on a 10-day course of tamoxifen treatment. Although robust expression of the ZsGreen was observed in the liver (Fig. 1 A), no ZsGreen fluorescence was appreciated in the bone marrow (Fig. 1 B). Within the liver, ZsGreen were from HNF4a^+^ hepatocytes (Fig. 1 C and D). These data suggest that hepatocytes but not the bone marrow is the major source of TPO after 5-FU treatment.

**Figure 1.**
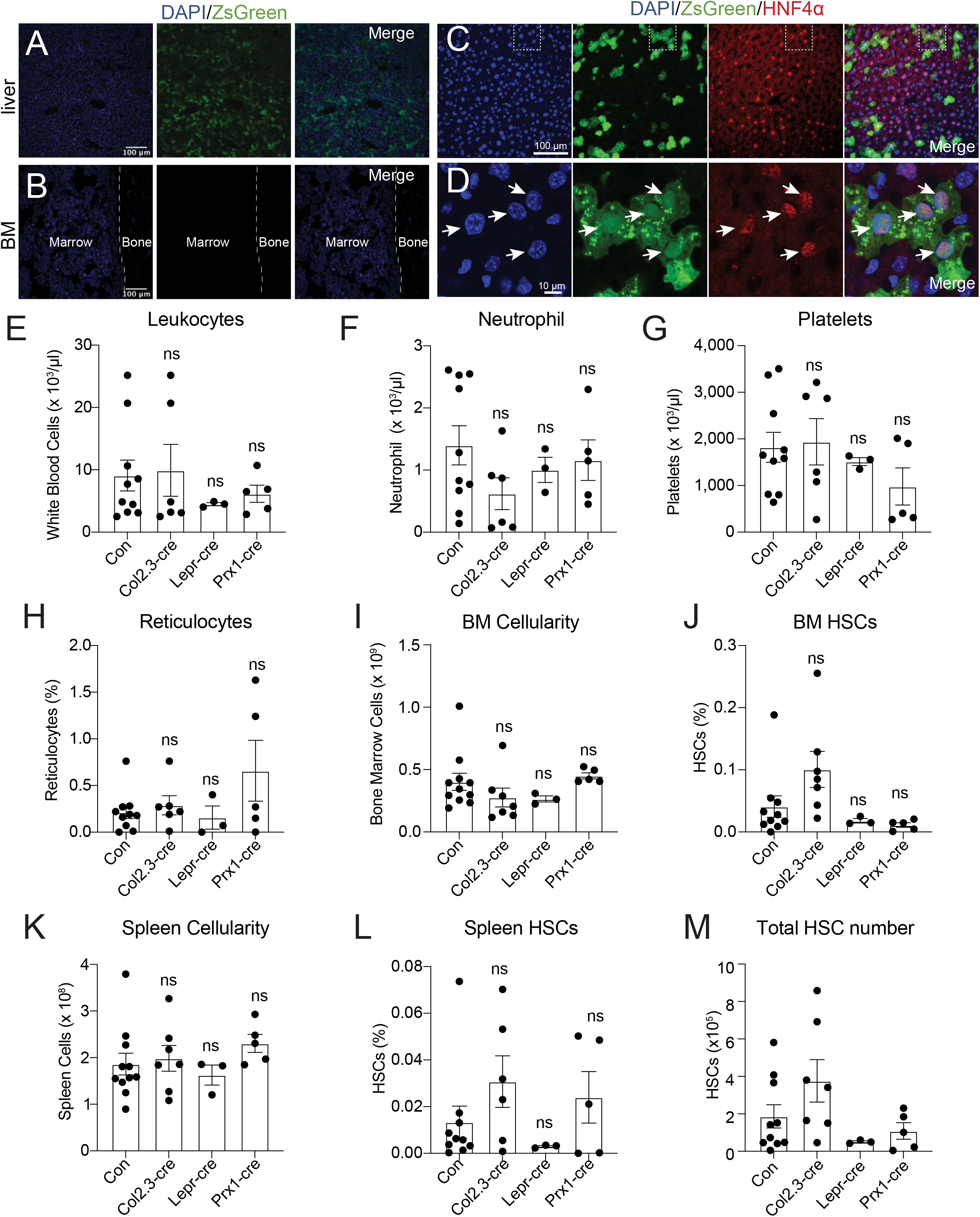
Bone marrow TPO is not required for hematopoietic recovery and HSC expansion after 5-FU chemoablation. (A) Confocal images of liver sections from tamoxifen-treated *Tpo^creER^; loxpZsGreen* mice 10 days after 5-FU injection. (B) Confocal images of femur sections from tamoxifen-treated *Tpo^creER^; loxpZsGreen* mice 10 days after 5-FU injection. (C and D) Confocal images showing immunostaining of HNF4α on liver sections from tamoxifen-treated *Tpo^creER^; loxpZsGreen* mice 10 days after 5-FU injection. (E-H) Blood counts of mice with *Tpo* conditionally deleted from osteoblasts, bone marrow stromal cells, or both, and controls after 5-FU challenge. n=3-10 mice. (I-L) Cellularity and frequencies of HSCs in the bone marrow and spleens from mice with *Tpo* conditionally deleted from osteoblasts, bone marrow stromal cells, or both, and controls after 5-FU challenge. n=3-10 mice. (M) Total number of HSCs in the bone marrow and spleens from mice with *Tpo* conditionally deleted from osteoblasts, bone marrow stromal cells, or both, and controls after 5-FU challenge. n=3-10 mice. Con: *Tpo^gfp/+^* control mice. Col2.3-cre: *Col2.3-cre; Tpo^fl/gfp^* mice. Lepr-cre: *Lepr-cre; Tpo^fl/gfp^* mice. Prx1-cre: *Prx1-cre; Tpo^fl/gfp^* mice. Data represent mean ± sem. ns, not significant. Statistical significance was calculated with one-way ANOVA. Each dot represents one independent mouse in E-M.

Although we did not detect appreciable *Tpo* translation in the bone marrow after 5-FU treatment (Fig. 1 B), osteoblasts and LepR^+^ mesenchymal stromal cells are the major bone marrow cells expressing *Tpo* transcripts ^17,33,36^, and based on antibody staining, it has been reported that osteoblasts express TPO protein after 5-FU treatment ^19^. To formally test the function of TPO from these cells, we conditionally deleted *Tpo* from osteoblasts (*Col2.3-cre; Tpo^fl/gfp^*) or mesenchymal stromal cells (*Lepr-cre; Tpo^fl/gfp^*). Deletion of *Tpo* from osteoblasts or LepR^+^ stromal cells had no significant effects on leukocytes, neutrophils, platelets, reticulocytes, bone marrow and spleen cellularity, HSCs, or LSKs after 5-FU treatment (Fig. 1 E-M, Fig. S2 B and C). These results show that hematopoietic recovery and HSC regeneration from 5-FU treatment does not require TPO production by bone marrow osteoblasts or mesenchymal stromal cells.

It is possible that osteoblasts and mesenchymal stromal cells serve as redundant sources of TPO. We thus also generated *Prx1-cre; Tpo^fl/gfp^* mice. *Prx1-cre* drives recombination in mesenchymal lineage cells in murine long bones, including both osteoblasts and mesenchymal stromal cells ^3,37,38^, allowing conditional deletion of *Tpo* from both osteoblasts and mesenchymal stromal cells. Adult baseline *Prx1-cre; Tpo^fl/gfp^* mice had normal blood cell counts, normal bone marrow cellularity, normal HSC and LSK frequencies, and showed no signs of extramedullary hematopoiesis in the spleen (Fig. S2 D-N), consistent with the notion that TPO from the bone marrow is not required for HSC maintenance or hematopoiesis ^17^. We then administrated 5-FU to these mice. Deletion of *Tpo* from both osteoblasts and mesenchymal cells in *Prx1-cre; Tpo^fl/gfp^* mice had no effects on leukocytes, neutrophils, platelets, reticulocytes, bone marrow and spleen cellularity, HSCs, or LSKs after 5-FU chemoablation (Fig. 1 E-M, Fig. S2 B and C). These results strengthen our finding that the bone marrow is not a source of TPO for hematopoietic recovery and HSC regeneration following 5-FU treatment.

Hepatic TPO is required for steady-state bone marrow HSC maintenance as conditional deletion of *Tpo* from hepatocytes leads to HSC depletion ^17^. We found that hepatocytes are also the major source of TPO after 5-FU treatment (Fig. 1 A-D). To address whether hepatic TPO is required for hematopoietic recovery and HSC regeneration after 5-FU-mediated stress, we treated adult *Tpo^fl/fl^* mice with hepatotropic AAV8-TBG-cre virus (AAV) concurrently with 5-FU injection. A single dose of this *Cre*-bearing adenovirus drives efficient and specific recombination in hepatocytes ^17^. Ten days after the treatment, AAV-treated mice showed a significant reduction of platelet and reticulocyte expansion, with normal leukocyte and neutrophil responses relative to controls (Fig. 2 A-D, Fig. S3 A and B). Although the response of spleen HSC frequency to 5-FU was not significantly impacted by AAV treatment, the responses of bone marrow HSC frequency and the total number of HSCs were significantly decreased in AAV-treated mice compared to controls (Fig. 2 E-N), indicating a compromised HSC recovery after 5-FU treatment. Consistently, bone marrow cells from these AAV-treated mice after 5-FU administration showed a significant reduction in reconstituting irradiated recipient mice compared with controls (Fig. 2 O). The responses of bone marrow and spleen LSK frequencies to 5-FU were largely normal in AAV-treated mice compared with controls (Fig. S3 C and D), suggesting that the effects of acutely disrupted TPO signaling are most pronounced on HSCs, rather than restricted hematopoietic progenitors. Altogether, these data suggest that hepatic but not bone marrow-derived TPO is required for hematopoietic and HSC recovery after 5-FU treatment.

**Figure 2.** Hepatic TPO is required for hematopoietic recovery and HSC expansion after 5-FU chemoablation. (A-D) Platelet counts and reticulocyte frequencies (A and C) and normalized fold change relative to no treatment baseline (B and D) of mice with *Tpo* conditionally deleted from hepatocytes and their respective controls with and without 5-FU challenge. n=4-16 mice. (E-L) Cellularity and frequencies of HSCs (E, G, I, K) and normalized fold change (F, H, J, L) in the bone marrow and spleens from mice with *Tpo* conditionally deleted from hepatocytes and controls with and without 5-FU challenge. n=4-16 mice. (M and N) Total number of HSCs (M) and normalized fold change (N) in the bone marrow and spleens from mice with *Tpo* conditionally deleted from hepatocytes and controls with and without 5-FU challenge. n=4-16 mice. (O) Numbers of total recipients and long-term multilineage reconstituted recipients after transplantation of 50,000 bone marrow cells from PBS- or AAV-treated *Tpo^fl/fl^* and 5-FU challenged mice along with 500,000 unchallenged competitor bone marrow cells. Recipients were scored as reconstituted if they showed donor-derived myeloid, B and T cells in the peripheral blood 16 weeks after transplantation. Data were from recipients of three independent donor pairs from two independent experiments. Con: *Tpo^fl/fl^* or wild-type control mice. AAV: AAV8-TBG-cre-treated *Tpo^fl/fl^* mice. Data represent mean ± sem. ns, not significant, *P<0.05, ** P<0.01, *** P<0.001, ****P<0.0001. Statistical significance was assessed with two-way ANOVA (A, C, E, G, I, K, M), student’s t-test (B, D, F, H, J, L, N), or Fisher’s exact test (O). Each dot represents one independent mouse in A-N.

### Hematopoiesis from TPO-deficient mice are sensitive to ionizing radiation

Ionizing radiation is another common myeloablative treatment. To test whether TPO is required for hematopoietic recovery in this condition, we exposed mice to a single 5.5 Gy dose of radiation. Four weeks after the irradiation, both wild-type and *Tpo^gfp/gfp^* mice had leukopenia, decreased bone marrow cellularity, and decreased bone marrow LSKs relative to non-irradiated baseline levels (Fig. S4 A-F). However, *Tpo^gfp/gfp^* mice had significantly more severe perturbations and additionally had no detectable bone marrow HSCs (Fig. S4 A-G). The functional impact of these differences was demonstrated by a significant increase in mortality among *Tpo^gfp/gfp^* mice following irradiation (Fig. S4 H). Compared with baseline levels, neither wild-type nor *Tpo^gfp/gfp^* mice had notable differences in platelet counts, spleen cellularity, or spleen LSK frequency in response to irradiation (Fig. S4 C, I and J). However, *Tpo^gfp/gfp^* mice had diminished regeneration of spleen HSC frequency and total HSC numbers compared with controls (Fig. S4 K and L). Thus, *Tpo* is required for hematopoietic and HSC recovery after irradiation.

### Hepatic but not bone marrow TPO is required HSC regeneration after ionizing radiation

It is not clear what cells are the major source of TPO after irradiation. Bone marrow stromal cells (CD45^-^ Ter119^-^) did not have significantly upregulated *Tpo* transcripts 14 days after irradiation (Fig. S2 A). To assess the translation of TPO protein following irradiation, *Tpo^creER^; ZsGreen^LSL^* translational reporter mice were irradiated and immediately dosed with a 10-day course of tamoxifen treatment. Although robust recombination of the *ZsGreen^LSL^* allele was observed in the liver (Fig. 3 A), no ZsGreen was observed in the bone marrow (Fig. 3 B). Hepatocytes were the major cell type that expressed TPO in the liver after irradiation (Fig. 3 C and D). This finding suggests that hepatocytes are the major source of TPO with no appreciable expression in the bone marrow following radioablation.

**Figure 3.**
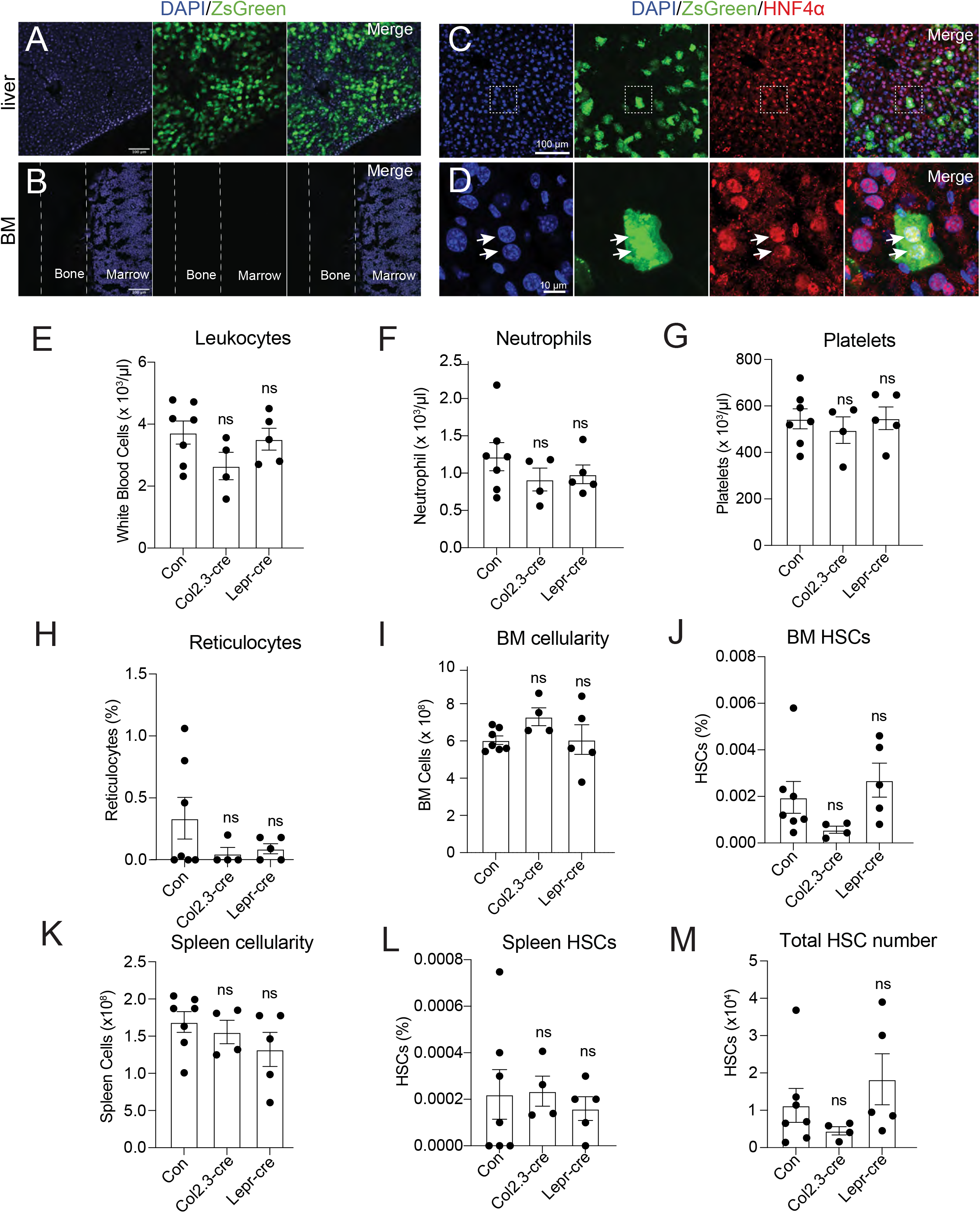
Bone marrow TPO is not required for hematopoietic recovery from irradiation. (A) Confocal images of liver sections from tamoxifen-treated *Tpo^creER^; loxpZsGreen* mice 10 days after irradiation. (B) Confocal images of femur sections from tamoxifen-treated *Tpo^creER^; loxpZsGreen* mice 10 days after irradiation. (C and D) Confocal images showing immunostaining of HNF4α on liver sections from tamoxifen-treated *Tpo^creER^; loxpZsGreen* mice 10 days after irradiation. (E-H) Blood counts of mice with *Tpo* conditionally deleted from osteoblasts, bone marrow stromal cells, and controls after irradiation. n=4-7 mice. (I-L) Cellularity and frequencies of HSCs in the bone marrow and spleens from mice with *Tpo* conditionally deleted from osteoblasts, bone marrow stromal cells, and controls after irradiation. n=4-7 mice. (M) Total number of HSCs in the bone marrow and spleens from mice with *Tpo* conditionally deleted from osteoblasts, bone marrow stromal cells, and controls after irradiation. n=4-7 mice. Con: *Tpo^gfp/+^* control mice. Col2.3-cre: *Col2.3-cre; Tpo^fl/gfp^* mice. Lepr-cre: *Lepr-cre; Tpo^fl/gfp^* mice. Prx1-cre: *Prx1-cre; Tpo^fl/gfp^* mice. Data represent mean ± sem. ns, not significant. Statistical significance was assessed with one-way ANOVA. Each dot represents one independent mouse in E-M.

To functionally test whether osteoblasts and bone marrow mesenchymal cells are sources of TPO for hematopoietic recovery and HSC regeneration after irradiation, we gave *Col2.3-cre; Tpo^fl/gfp^* or *Lepr-cre; Tpo^fl/gfp^* mice, along with littermate controls, a single 5.5 Gy dose of radiation and analyzed them four weeks later. Compared with irradiated controls, *Col2.3-cre; Tpo^fl/gfp^*, or *Lepr-cre; Tpo^fl/gfp^*, mice showed no significant differences in leukocytes, neutrophils, platelets, and reticulocytes (Fig. 3 E-H). Bone marrow cellularity, LSK and HSC frequencies were also normal (Fig. 3 I and J, Fig. S5 A). These mice also had normal spleen cellularity, LSK and HSC frequency, as well as total HSC numbers (Fig. 3 K-M and Fig. S5 B). These data suggest that osteoblasts or the bone marrow mesenchymal stromal cells are a dispensable source of TPO for HSC and hematopoietic recovery from radioablative challenges.

To test the role of hepatocyte-derived TPO, we then irradiated *Tpo^fl/fl^* mice treated with AAV8-TBG-cre virus. Compared to control mice, these mice showed a similar response to irradiation in leukocyte, neutrophil and platelet counts as well as reticulocyte frequency (Fig. 4 A, and Fig. S5 C-E). Although the bone marrow cellularity response to irradiation was similar between *Tpo^fl/fl^* mice treated with AAV and control, there was a significant reduction in bone marrow HSC but not LSK frequency recovery relative to controls after irradiation (Fig. 4 C-F and Fig. S5 F). Compared with controls, spleen cellularity, LSK and HSC frequencies from *Tpo^fl/fl^* mice treated with AAV had a similar recovery after irradiation (Fig. 4 G-J and Fig. S5 G). Overall, *Tpo^fl/fl^* mice treated with AAV had a significant reduction in total HSC number recovery after irradiation (Fig. 4 K and L). Importantly, bone marrow cells from these irradiated AAV-treated mice showed functional defects in reconstituting irradiated recipients (Fig. 4 M). These data suggest that hepatic TPO is required for proper functional recovery of HSCs following irradiation-mediated myeloablation.

**Figure 4.**
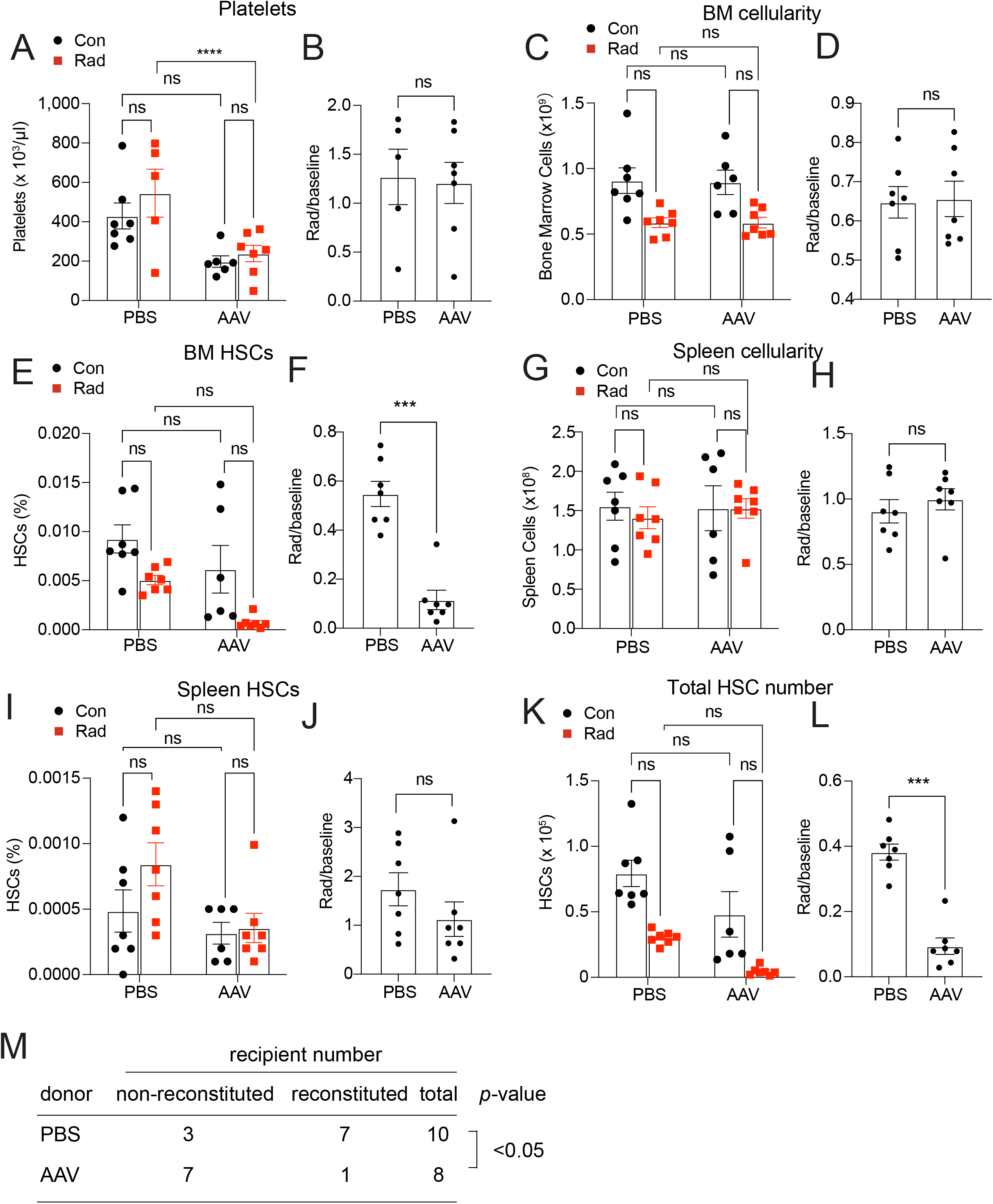
Hepatic TPO is required for effective HSC recovery from ionizing radiation. (A and B) Platelet counts (A) and normalized fold change (B) in mice with *Tpo* conditionally deleted from hepatocytes and controls with and without irradiation. n=5-7 mice. (C-J) Cellularity and frequencies of HSCs in the bone marrow and spleens from mice with *Tpo* conditionally deleted from hepatocytes and controls with and without irradiation. n=6-7 mice. (K and L) Total number of HSCs in the bone marrow and spleens from mice with *Tpo* conditionally deleted from hepatocytes with and without irradiation. n=6-7 mice. (M) Numbers of total recipients and long-term multilineage reconstituted recipients after transplantation of 1,000,000 bone marrow cells from PBS- or AAV-treated *Tpo^fl/fl^* and irradiated donor mice along with 1,000,000 irradiated competitor bone marrow cells. Recipients were scored as reconstituted if they showed donor-derived myeloid, B and T cells in the peripheral blood 16 weeks after transplantation. Data were from independent recipients of two donor pairs from two independent experiments. Con: *Tpo^fl/fl^* or wild-type control mice. AAV: AAV8-TBG-cre-treated *Tpo^fl/fl^* mice. Data represent mean ± sem. ns, not significant, *** P<0.001. Statistical significance was assessed with two-way ANOVA (A, C, E, G, I, K), student’s t-test (B, D, F, H, J, L), or Fisher’s exact test (M). Each dot represents one independent mouse in A-L.

## Discussion

Our data show that the bone marrow does not meaningfully upregulate TPO protein after 5-FU treatment or irradiation. Although we did not exhaustively investigate all models of ablative conditioning, it appears that the bone marrow is not a functional source of TPO in hematopoietic recovery from acute myeloablative stress. However, our data do not rule out the possibility that bone marrow may serve as a source of TPO in certain specific stress conditions, such as idiopathic thrombocytopenia purpura ^33^. Future investigation into additional stress conditions is needed. We found that liver hepatocytes are an important source of TPO for stress hematopoiesis. Although several other studies have investigated the role of TPO on hematopoietic recovery after stress, the source of TPO and its impact on HSCs (rather than hematopoietic progenitors) have not been examined. By examining HSCs and hematopoietic progenitors, we show that HSCs are more sensitive to acute TPO perturbation than hematopoietic progenitors (LSKs), suggesting the major role of TPO in myeloablation is to promote HSC regeneration, at least under the conditions we studied. To our knowledge, TPO is the first systemic factor required for HSC regeneration and hematopoietic recovery identified to date. The presence of a stem-cell hormone, like TPO, raises the possibility of other endocrine regulators of HSC regeneration and hematopoietic recovery.

Numerous connections between hepatic and hematopoietic pathophysiology have long been appreciated. Aplastic anemia is a known sequela of pediatric liver failure, and thrombocytopenia is a well-characterized feature of hepatic disease ^39–41^. While mechanisms such as metabolic toxicity and splenic sequestration may play a role in mediating these effects, it seems likely that disruption of TPO signaling is also involved. Indeed, several recent clinical studies have shown that newly developed TPO agonists can reduce the need for platelet transfusions in patients with chronic liver disease ^42,43^. While early trials of TPO mimetics in primary and acquired aplastic anemia have been promising, to our knowledge, no group has specifically studied their use in liver failure-associated aplastic anemia ^31,32^. Given that hepatic *Tpo* mRNA and serum TPO levels are significantly downregulated in liver failure ^44^, further studies are warranted. Our data suggest that cancer patients with underlying liver disease may have greater sensitivity to myeloablative conditioning and supplementing TPO mimetics may help alleviate the adverse effects associated with myeloablative conditioning in these patients.

## Materials and methods

### Mice

All mice were 8-16 weeks old and maintained on a C57BL/6 background. *Prx1-cre, Lepr-cre* and *LoxpZsGreen* mice were obtained from the Jackson Laboratory. *Col2.3-cre* mice were described previously ^45^. Generation of *Tpo^gfp^, Tpo^creER^* and *Tpo^fl^* mice was described previously ^17^. All mice were housed in specific pathogen-free, Association for the Assessment and Accreditation of Laboratory Animal Care (AAALAC)-approved facilities at the Columbia University Medical Center. All protocols were approved by the Institute Animal Care and Use Committee of Columbia University.

### Genotyping PCR

Primers for genotyping *Tpo^creER^*: OLD815, 5’-CCACCACCATGCCTAACTCT-3’; OLD816, 5’-GTTCTCCTCCACGTCTCCAG-3’; and OLD817, 5’-TCGCTAGCTGCTCTGATGAA-3’.

Primers for genotyping *loxpZsGreen*: GGCATTAAAGCAGCGTATCC and AACCAGAAGTGGCACCTGAC

Primers for genotyping *Tpo^fl^*: OLD581, 5’-CATCTCGCTGCTCTTAGCAGGG-3’ and OLD582, 5’-GAGCTGTTTGTGTTCCAACTGG-3’.

Primers for genotyping *Tpo^gfp^*: OLD292, 5’-CGGACACGCTGAACTTGTGG-3’; OLD528 5’-ACTTATTCTCAGGTGGTGACTC-3’ and OLD653 5’-AGGGAGCCACTTCAGTTAGAC-3’.

Primers for genotyping *Lepr-cre:* OLD434 5’-CATTGTATGGGATCTGATCTGG-3’ and OLD435 5’-GGCAAATTTTGGTGTACGGTC-3’.

Primers for genotyping *cre*: OLD338, 5’-GCATTTCTGGGGATTGCTTA-3’ and OLD339, 5’-ATTCTCCCACCGTCAGTACG-3’.

### Chemoablative challenge

5-Fluorouracil (5-FU) (150mg/kg) was administered to mice *via* a single intraperitoneal injection. Ten days post-injection, mice were euthanized for analysis.

### Radioablative challenge

Mice were irradiated by a Cesium 137 Irradiator (JL Shepherd and Associates) at 300 rad/min with one dose of 550 rad. Four weeks after irradiation, mice were euthanized for analysis.

### Tamoxifen administration

Tamoxifen (Sigma) was dissolved in corn oil for a final concentration of 20mg/mL. Every other day for 10 days, 50uL of the solution was administered by oral gavage. Mice were analyzed 2-4 days after the final tamoxifen administration.

### Viral infections

Replication-incompetent AAV8-TBG-cre was obtained from the Penn Vector Core or Addgene. AAV8-TBG-cre carries Cre recombinase under the regulatory control of hepatocyte-specific thyroid-binding globulin (TBG) promoter. Efficient recombination was achieved at a dose of 2.5 x 10^11^ viral particles diluted in sterile 1x PBS. AAV8-TBG-cre was administered to mice *via* retro-orbital venous sinus injection.

### Flow cytometry

Bone marrow cells were isolated by flushing the long bones or by crushing the long bones, pelvis, and vertebrae with mortar and pestle in Ca^2+^ and Mg^2+^free HBSS with 2% heat-inactivated bovine serum. Spleen cells were obtained by crushing the spleens between two glass slides. The cells were passed through a 25G needle several times and filtered with a 70μm nylon mesh. The following antibodies were used to perform HSC staining: lineage markers (anti-CD2 (RM2-5), anti-CD3 (17A2), anti-CD5 (53-7.3), anti-CD8a (53-6.7), anti-B220 (6B2), anti-Gr-1 (8C5), anti-Ter119), anti-Sca-1 (E13-161.7), anti-c-kit (2B8), anti-CD48 (HM48-1), anti-CD150 (TC15-12F12.2).

For flow cytometric sorting of bone marrow stromal cells, bone marrow plugs were flushed as described above, then digested with collagenase IV (200 U/mL) and DNase1 (200 U/mL) at 37°C for 20min. Samples were then stained with anti-CD45 (30F-11), and anti-Ter119 antibodies and sorted on a flow cytometer.

### qRT-PCR

Cells were sorted directly into Trizol. Total RNA was extracted according to manufacturer’s instructions. Total RNA was subjected to reverse transcription using SuperScript III (Invitrogen) or ProtoScript II (NEB). Quantitative real-time PCR was run using SYBR green on a CFX Connect system (Biorad). β-actin was used to normalize the RNA content of samples. Primers used were: *Tpo*: OLD390, CCTTTGTCTATCCCTGTTCTGC and OLD391, ACTGCCCCTAGAATGTCCTGT. *β-actin:* OLD27, 5’-GCTCTTTTCCAGCCTTCCTT-3’ and OLD28, 5’-CTTCTGCATCCTGTCAGCAA-3’.

### Bone and liver sectioning

Freshly disassociated long bones were fixed for 3h in a solution of 4% paraformaldehyde, 7% picric acid, and 10% sucrose (W/V). The bones were then embedded in 8% gelatin, immediately snap frozen in liquid N_2_, and stored at −80 °C. Bones were sectioned using a CryoJane system (Instrumedics). For liver, cardiac perfusion with formalin was performed immediately after mouse sacrifice, and perfused liver tissue was dehydrated in a 30% sucrose solution overnight at 4 °C. Liver tissue was then placed in PELCO Cryo-Embedding compound (Ted Pella, Inc.), frozen on dry ice, and stored at −80 °C. Liver tissue was sectioned and directly transferred onto microscope slides. Both bone and liver sections were dried overnight at room temperature and stored at −80 °C. Sections were rehydrated in PBS for 5min, stained with DAPI for 15min, then mounted with Vectashield Hardset Antifade Mounting Medium (Vector Laboratories). Images were acquired on a Nikon Eclipse Ti confocal microscope (Nikon Instruments) or a Leica SP8 confocal microscope (Leica Microsystems).

### Long-term competitive reconstitution assay

Adult recipient mice were lethally irradiated by a Cesium 137 Irradiator (JL Shepherd and Associates) at 300 rad/min with two doses of 550 rad (total 1100 rad) delivered at least two hours apart. Cells were transplanted by retro-orbital venous sinus injection of anesthetized mice. Donor bone marrow cells were transplanted along with recipient bone marrow cells into lethally irradiated recipient mice. The recipient bone marrow cells were from mice that are treated similarly as the donor mice. Mice were maintained on antibiotic water (Baytril 0.17g/L) for 14 days then switched to regular water. Recipient mice were periodically bled to assess the level of donor-derived blood lineages, including myeloid, B, and T cells for at least 16 weeks. Blood was subjected to ammonium chloride potassium red cell lysis before antibody staining. Antibodies including anti-CD45.2 (104), anti-CD45.1 (A20), anti-CD3 (17A2), anti-B220 (6B2), anti-Gr-1 (8C5), and anti-Mac-1 (M1/70) were used to stain cells. Mice with presence of donor-derived myeloid, B, and T cells for 16 weeks were considered as long-term multilineage reconstituted.

### Statistical Analysis

All analyses were done using GraphPad Prism 7.0. In any comparison between a pooled control cohort and multiple experimental conditions, we used one-way ANOVA with Dunnett’s test. For all other comparisons we used unpaired Welch’s t-test. P-values and adjusted p-values between 0.01 and 0.1 were displayed; those greater than 0.1 were designated ‘not significant’. In all figures, error bars represent standard error of mean.

### Data Sharing Statement

For original data, please contact the corresponding author by email.

## Acknowledgements

This work was supported by the National Heart, Lung and Blood Institute (R01HL132074). L.G. was supported by the NYSTEM Columbia training program in stem cell research and a Columbia Stem Cell Initiative Seed Grant. M.D. was supported by the Columbia Medical Scientist Training Program and the NIH (1F30HL137323). L.D. was supported by the Rita Allen Foundation, the Irma Hirschl Research Award, the Schaefer Research Scholar Program and the Leukemia and Lymphoma Society Scholar award, R01HL153487 and R01HL155868. Images were collected in the Confocal and Specialized Microscopy Shared Resource of the Herbert Irving Comprehensive Cancer Center at Columbia University, supported by NIH grant P30CA013696 (National Cancer Institute). We thank M. Kissner at the Columbia Stem Cell Initiative for help on flow cytometry.

## Author Contributions

L.G., M.D., and H.C. performed all of the experiments. L.G., M.D., and L.D. designed the experiments, interpreted the results, and wrote the manuscript.

## Author Information

The authors declare no competing financial interests.

---

**FIGURE S1.**
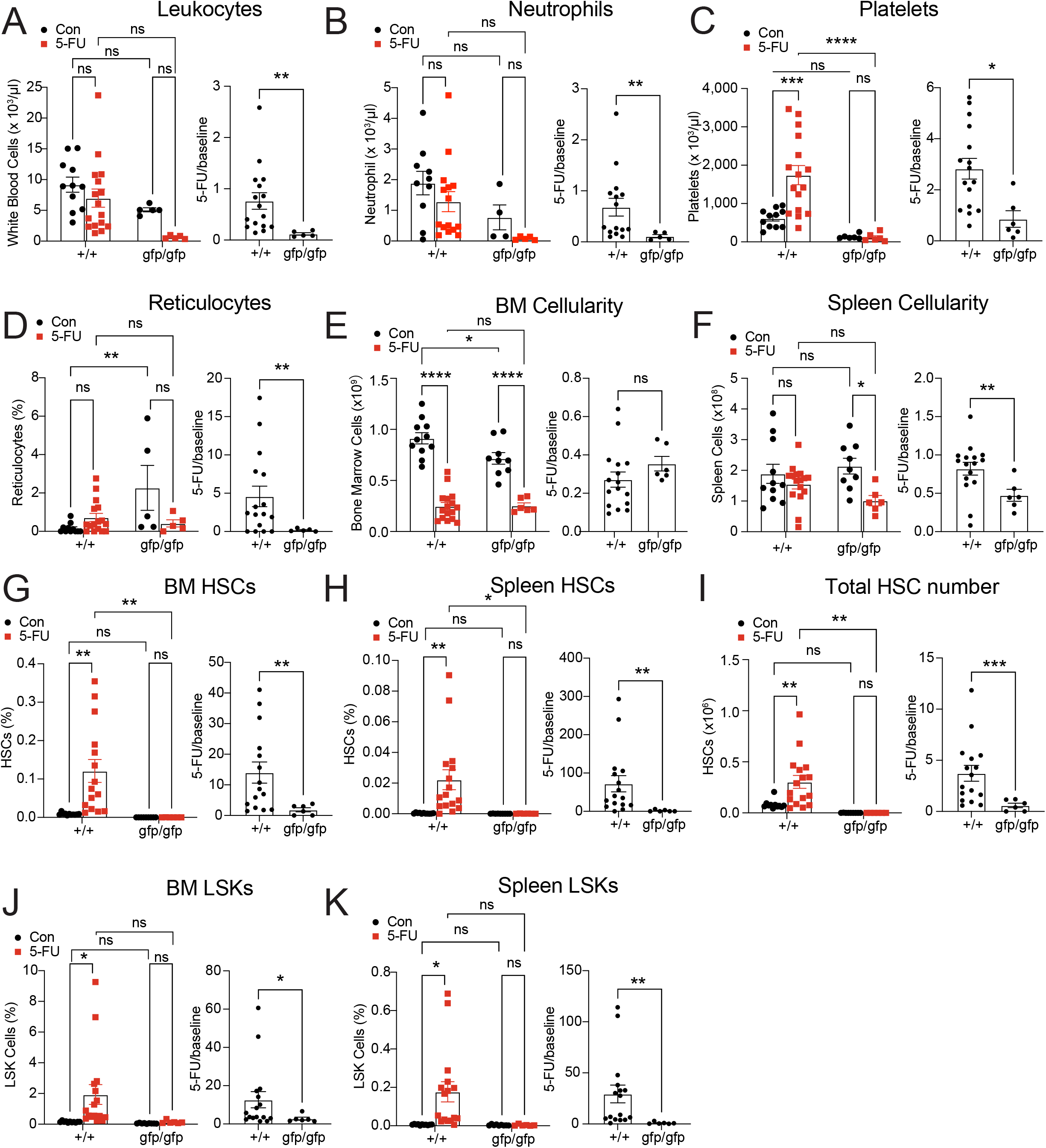

**FIGURE S2.**
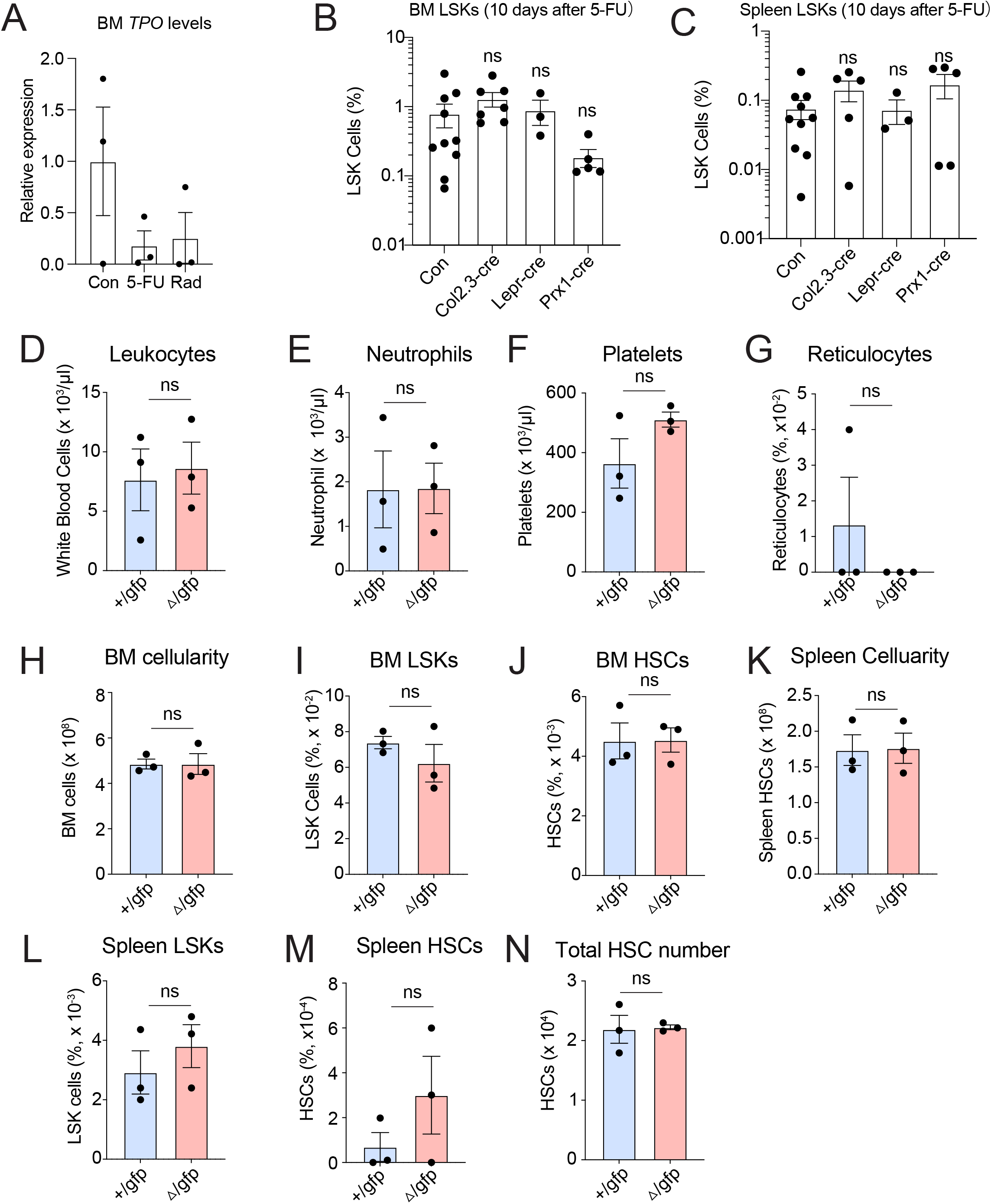

**FIGURE S3.** 

**FIGURE S4.**
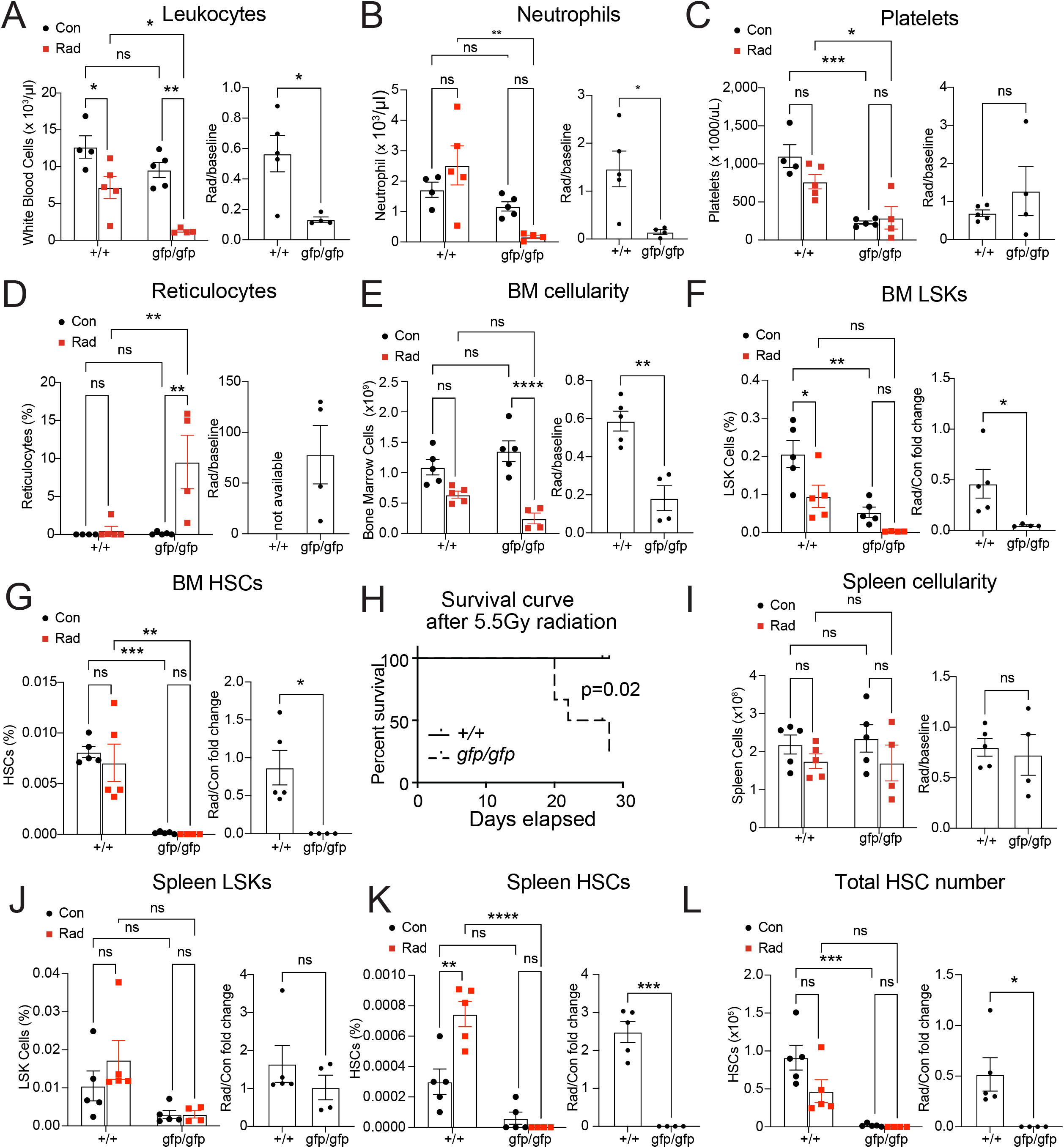

**FIGURE S5.**
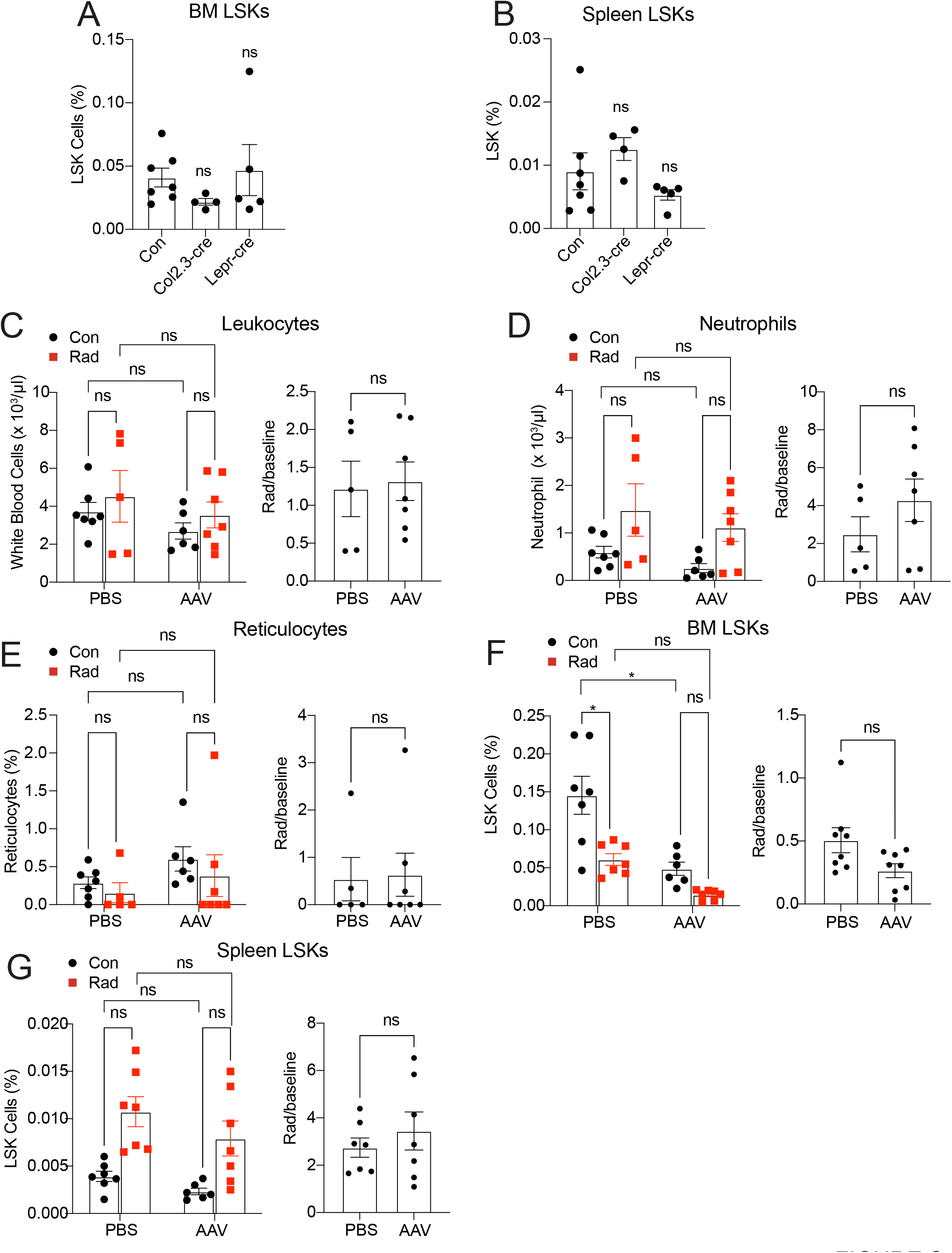

## Notes

### Competing Interest Statement

The authors have declared no competing interest.

